# Hippocampal CA2 Inhibition Disrupts Prefrontal and Thalamic Connectivity

**DOI:** 10.64898/2026.08.25.746953

**Authors:** Alessa A. Franz, Tudor M. Ionescu, Dennis Kätzel, Bastian Hengerer

## Abstract

Disturbances in the CA2-subfield of the hippocampus have been associated with symptoms of psychiatric disorders, including impaired social behavior. Using chemogenetic inhibition during functional ultrasound imaging, we found that dorsal CA2 pyramidal neurons broadly control prefrontal and thalamic communication, in addition to hippocampal and thalamic activity. Correspondingly, chronic CA2 inhibition altered social interaction.

## Main text

Neuropsychiatric disorders such as schizophrenia are characterized by impaired social functioning^1^. However, the underlying causes are still unclear and effective pharmacological treatments targeting social cognition are still lacking^1,2^. The hippocampal subregion CA2 has recently been identified as critical to social memory in rodents^3^, as intact excitatory CA2 output^3,4^ as well as some extrahippocampal input to CA2^5^ are crucial for social recognition memory in mice. Preclinical^6^ and clinical^7^ data have associated disturbances in the neuronal integrity of CA2 with psychiatric pathologies. Hence, understanding CA2’s relevance in global network function and behavioral control is important for providing further insights into the pathomechanism of social pathologies.

To mimic a pathological hypoexcitability of CA2 pyramidal neurons of the murine dorsal hippocampus (dHPC)^6^, we transduced Amigo2-Cre mice virally with the inhibitory chemogenetic receptor hM4Di. The selective targeting of dorsal CA2 pyramidal neurons was shown and the bioavailability of the hM4Di-agonist clozapine-N-oxide (CNO) verified by quantifying its metabolite clozapine in plasma (Fig. S1). To identify the causal role of CA2 on brain-wide activity and connectivity, we used functional ultrasound (fUS) whole-brain imaging of cerebral blood volume (CBV)^8^ in hM4Di-transduced (hM4Di^+^) or mCherry-transduced control, sedated mice. fUS acquisitions before and after acute CNO injection (Fig. 1a) revealed a significant CNO-induced reduction in dCA2 activity of hM4Di^+^ mice relative to control (Fig. 1b), verifying the effective chemogenetic inhibition of dCA2 at the physiological level. Interestingly, CA2 inhibition also caused reduced activity in the CA1 subregions of the dHPC (dCA1) as well as the CA3 subfield of the ventral hippocampus (vHPC) (Fig. 1c, Fig. S2a, b). Despite CA2’s impact on activity levels, we observed no significant changes in the functional connectivity (FC) within the hippocampal network except for a left-hemispheric increase in ipsilateral FC between dCA2 and dCA3 in hM4Di^+^ mice (Fig.1f).

**Fig. 1.**
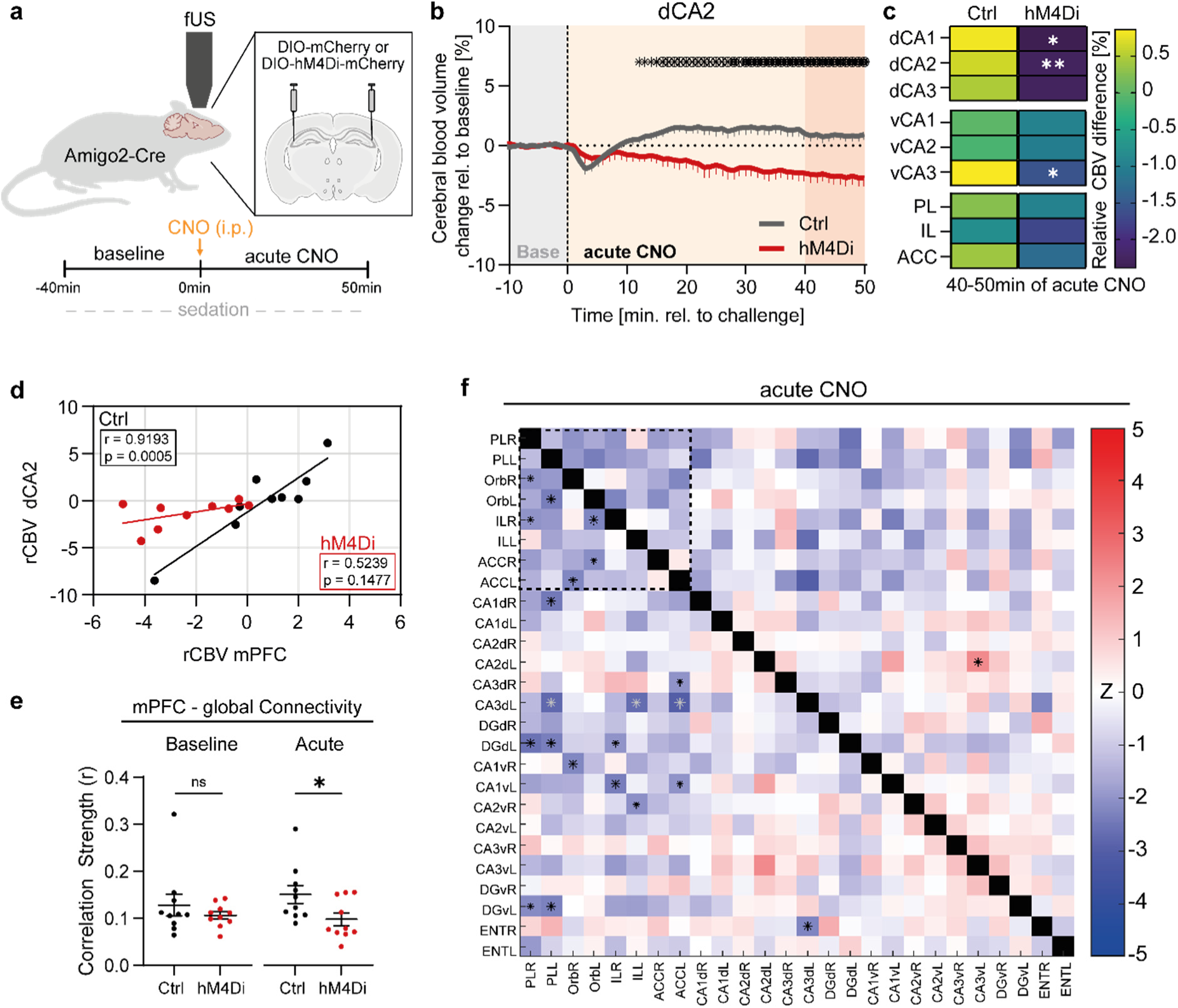
CA2 inhibition disrupts prefrontal connectivity. **a** Experimental setup of functional Ultrasound measurements. **b** Time course of changes in Cerebral Blood Volume (CBV) level in dorsal CA2 during baseline recording and after acute CNO application. Dashed line indicates CNO application. Dark reddish area indicates the investigation period for regional activity (n = 9 mice per group; Ancova, Group mean ± SEM, asterix is p < 0.05, circled asterix is p < 0.01, dark asterix is p < 0.001). **c** Percentual mean difference in relative CBV (rCBV) of subregions of the dorsal hippocampus (upper), ventral hippocampus (middle) and prefrontal cortex (lower) at minute 40-50 of scan (indicated by reddish area in **a**) (raw data are shown in Fig. S2; 2-tailed unpaired t-test: for dCA1, t(16) = 2.476, p = 0.0248; for dCA2, t(16) = 3.286, p = 0.0047; for vCA3, t(16) = 2.755, p = 0.0141). **d** Pearson r correlation in relative CBV between dCA2 and mPFC (for Vh, n = 9 mice; for hM4Di^+^, n = 10 mice). Trend line is generated via simple linear regression. **e** Global functional connectivity of the mPFC shown as average correlation at baseline and acute CNO (minutes 15-50) (acute for Vh (n = 10 mice) and hM4Di^+^ (n = 10 mice), 2-tailed unpaired t-test: t(18) = 2.285, p = 0.035). **f** Heatmap of functional connectivity between regions of the hippocampus and frontal cortex in hM4Di^+^ mice relative to controls during acute CNO (minutes 55-90). Dashed square indicates the mPFC-cluster of hypoconnectivity. Statistical difference is shown via Z-scores. Z < 0 is control > hM4Di^+^ (blue color), Z > 0 is Control < hM4Di^+^ (red color). *black: p < 0.05, *grey: p < 0.01, *white: p < 0.001. Otherwise, ns = not significant, *p < 0.05, ** p < 0.01. Statistical details are shown for significant changes only, all details are provided in statistical details file.

However, CA2’s connectivity to extra-HPC regions^9^ prompted us to next explore the effects of dCA2 inhibition brain-wide. Due to its role in social behaviors^10^ and dCA2’s modulatory influence on its gamma oscillations^11^ , we initially investigated the mPFC. Whereas acute CA2 inhibition did not seem to alter the *activity* in mPFC subregions significantly (Fig. S2c), it profoundly influenced their *connectivity* with other brain regions. Changes in connectivity were already indicated by an abolished positive correlation between spontaneous neuronal activities in dCA2 and mPFC in hM4Di mice relative to control (Fig. 1d). More specifically, acute CA2 inhibition induced a global hypoconnectivity of mPFC, particularly of prelimbic (PL) and infralimbic (IL) cortex (Fig. 1e, S3a-c). This reduced prefrontal FC was most prominent in connections with other frontal regions and the hippocampus (Fig. 1f and Fig. S5a). Within the mPFC-HPC network, only the FC between vHPC and mPFC was reduced (Fig. 1f and Fig. S3e-f). A persistent hypoconnected frontal network also emerged after chronic CA2 inhibition for 13 days (Fig. S3I and Fig. S5b).

Unexpectedly, brain wide analysis revealed a particularly prominent influence of CA2- modulation on the thalamus, indicated by changes in absolute (Fig. 2a) and relative CBV after acute CNO application (Fig. S2d and Fig. S4b). While the connectivity between dCA2 and thalamus seemed to remain stable, as indicated by the unchanged activity correlation between both regions (Fig. 2b) and the intact FC (Fig. S4d and S4g), the correlations between thalamic activity and both vHPC and mPFC activity were largely abolished in hM4Di^+^ mice (Fig. 2c-d; Fig. S4c). However, acute CA2 inhibition caused no changes in the overall global thalamic connectivity (Fig. 2e), suggesting circuit-specific alterations.

**Fig. 2.**
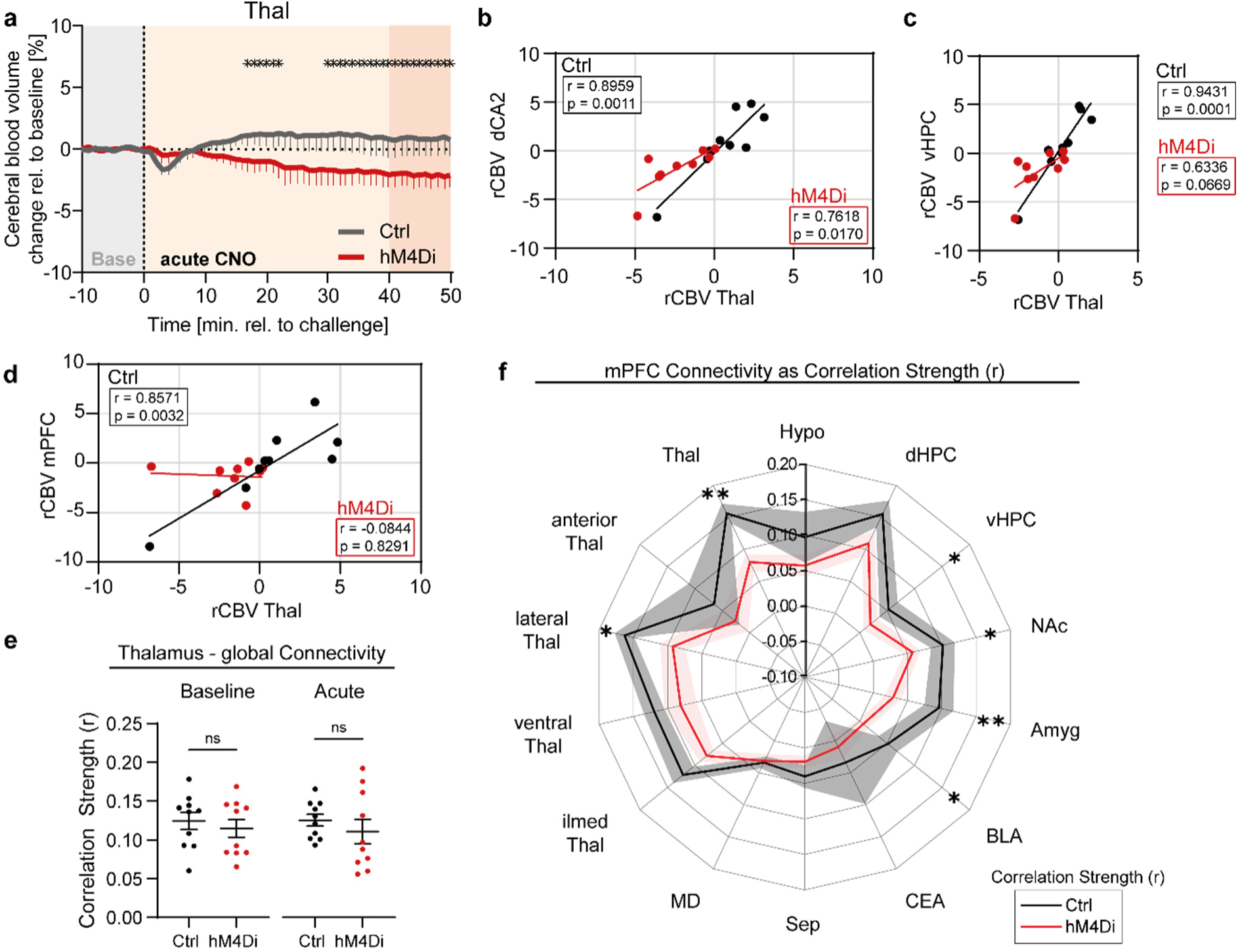
Disrupted thalamo-hippocampal and thalamo-mPFC connectivity. **a** Time course of changes in Cerebral Blood Volume (CBV) level in thalamus before and after acute CNO application. Dashed line indicates the timepoint of CNO treatment. Dark reddish area indicates the investigation period for regional activity (n = 9 mice per group; Ancova, Group mean ± SEM, asterix is p < 0.05). **b** Correlation in relative CBV between the dCA2 with the thalamus (for Vh, n = 9 mice; for hM4Di^+^, n = 10 mice). **c** Correlation in relative CBV of the ventral hippocampus with the thalamus (for Vh, n = 9 mice; for hM4Di^+^, n = 10 mice). **d** Correlation in relative CBV of the mPFC with the thalamus (for Vh, n = 9 mice; for hM4Di^+^, n = 10 mice). **e** Global functional connectivity of the thalamus shown as average correlation at baseline and acute CNO (minutes 15-50) (baseline for Vh, mean FC strength: 0.125 ± 0.011, n = 10 mice; for hM4Di^+^, mean FC strength: 0.115 ± 0.012, n = 10 mice; 2-tailed unpaired t-test: t(18) = 0.6055, p = 0.5524; acute for Vh, mean FC strength: 0.126 ± 0.008, n = 10 mice; for hM4Di^+^, mean FC strength: 0.112 ± 0.016, n = 10 mice; 2-tailed unpaired t-test: t(18) = 0.8281, p = 0.4184). **f** Spider plot of mPFC functional connectivity with brain regions following acute CNO shown as r-scores ± SEM (vHPC: 2-tailed unpaired Mann-Whitney t-test: U = 22, p = 0.0355; NAc: 2-tailed unpaired t-test: t(18) = 2.099, p = 0.0502; Amyg: 2-tailed unpaired Mann-Whitney t-test: U = 14, p = 0.0052; BLA: 2-tailed unpaired Mann-Whitney t-test: U = 17, p = 0.0115; Thal: for Vh, mean: 0.156 ± 0.016, for hM4Di, mean: 0.08 ± 0.013, 2- tailed unpaired t-test: t(18) = 3.773, p = 0.0014; lThal: 2-tailed unpaired Mann-Whitney t-test: U = 21, p = 0.0288). ilmed Thal is midline group, medial group and intralaminar nuclei of dorsal thalamus. Each data point represents a measured animal. Error bars are SEM. ns = not significant, *p < 0.05, ** p < 0.01. Trend line is generated via simple linear regression throughout. Statistical details are shown for significant changes only, all details are provided in statistical details file.

Subsequently, we assessed the frontal connectivity to the thalamus, a pathway recently associated with social memory ^12^ , additionally to other neuronal circuit nodes of social behavior^13^. Acute dCA2 inhibition reduced the connectivity of the mPFC with the thalamus – its lateral regions in particular – and with vHPC, nucleus accumbens and amygdala (Fig. 2f). Again, the FC of PL and IL, but not anterior cingulate cortex (ACC), with the thalamus was reduced (Fig. S4i). No changes were observed in overall dHPC- and vHPC-thalamic FC (Fig. S4e-f). In conclusion, dCA2 inhibition has wide-spread, but specific effects on the prefrontal and thalamic network.

Given these broad effects of dCA2-inhibition, especially on regions involved in social cognition, we next investigated its effects on the behavioral level. We aimed to emulate the chronic nature of psychiatric disease pathology^2,6^ by chronically inhibiting excitatory CA2 output, while monitoring social behavior of groups for extended periods (Fig. 3a). As mice are nocturnal animals, (6pm to 6am), we mostly concentrated on the dark phase (Fig. S6a-b). Automated video analysis^14^ revealed no effects of chronic CA2 inhibition on mobility, habituation, and the diurnal rhythm of social behavior (Fig. S6). Examining the social interaction with familiar mice (days 1 to 3), we observed that hM4Di^+^ mice exhibited more frequent short-term sniffing events with conspecifics, revealing a dark phase-specific subtle social interaction abnormality (Fig. 3b-d, Fig. S6e-g). To further investigate the social interaction and cognition of mice, we designed a test paradigm in which social interaction is measured on consecutive days over 22 h each. After spending the first four days in groups of familiar mice, two familiar mice were exchanged against 2 novel mice (day 5) and then changed back on day 6 (familiar group) (Fig. 3a, e). Analyzing the first two hours of each daily exposure (which overlapped with the beginning of the dark phase) as the initial contact period, we observed significantly higher social (Fig. 3f) and locomotion scores (Fig. 3g) on day 5 compared to day 4, indicating social interaction in both groups. Interestingly, control mice exhibited significantly less social interaction with familiar mice on day 6 compared to day 5 (mixed group), indicating a memory for the familiar mice which renders them less interesting after the exposure to novel mice. This pattern was not observed in hM4Di^+^ mice. On the contrary, locomotor activity was reduced in both groups on day 6 compared to day 5, indicating that dCA2 inhibition-induced alterations of behavior were specific to social stimuli. Using ‘social sniffing’ as a readout for close interaction between individuals, we further investigated this phenomenon by calculating “day vs day”-difference scores (Fig. 3h-j). Confirming the previous results (Fig. 3f), difference scores between days 6 and 5 revealed that control mice showed significantly less interaction with familiar mice (day 5) than with the mixed cohort (day 6), whereas hM4Di^+^ mice showed no difference in interaction between the two days (Fig. 3h). In contrast, no group difference was observed in social interaction between days 4 and 5, further supporting the notion that sociability remains intact following CA2 inhibition^3^. Our test paradigm included a 24 h separation from the familiar groups between days 4 and 6. Again, the difference score between days 6 and 4 was significantly lower in controls than in hM4Di^+^ mice, indicating that hM4Di^+^ mice showed a greater increase in interaction with conspecifics after encountering the mixed cohort compared with controls. This effect was also prominent during the first 25 min of the post-separation interaction phase (Fig.3j right). These results suggest that chronic DREADD-mediated CA2 inhibition leads to altered social interaction in groups, characterized by intensified, repetitive approach-behavior and longer-lasting impairment in social recognition.

**Fig. 3.**
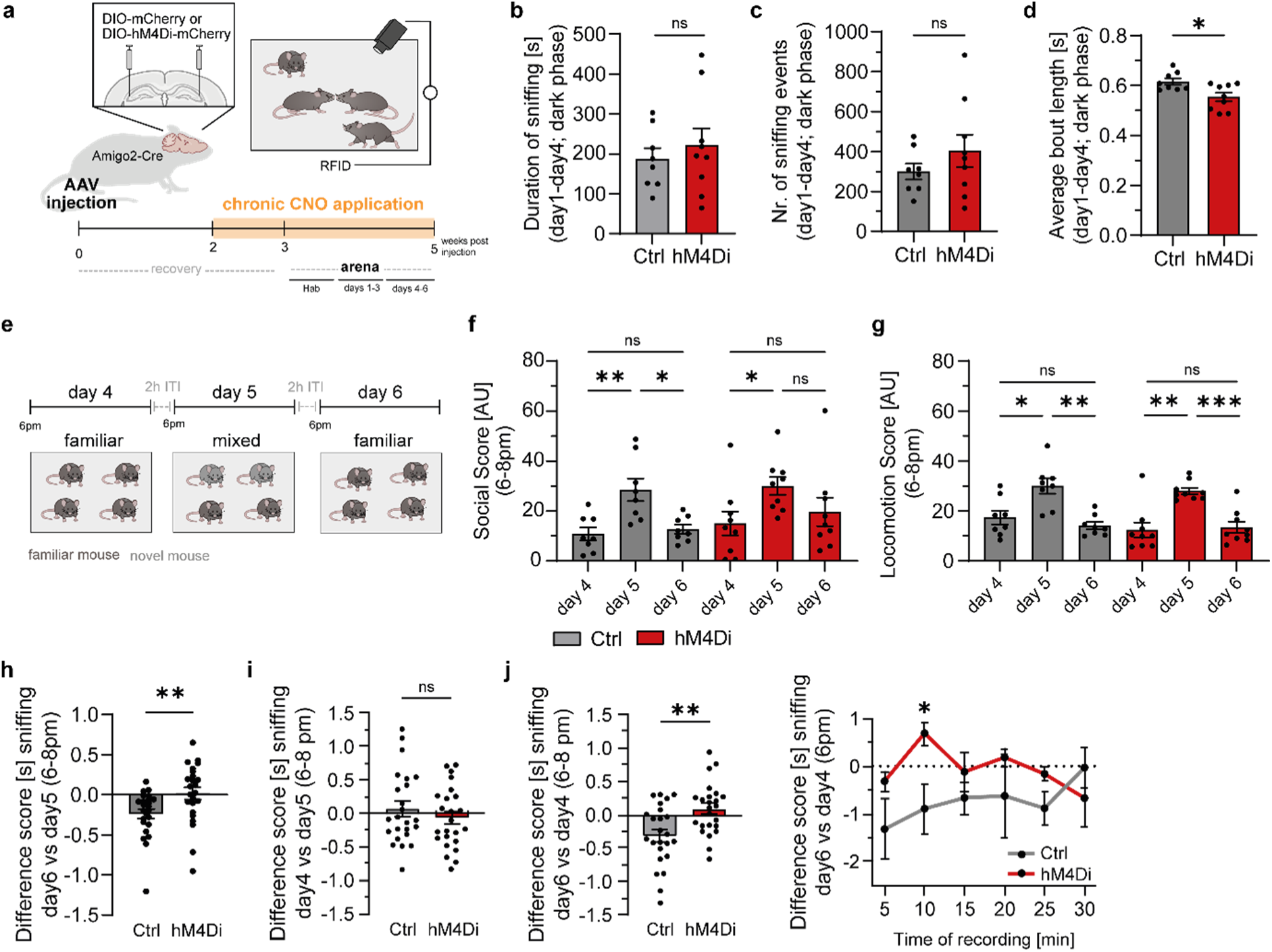
Altered nocturnal social interaction with familiar mice following chronic CA2 inhibition. **a** Timeline and experimental setup of behavioral testing. **b** Average of accumulated time mice spent sniffing with familiar mice during dark cycles (6pm-6am) of days 1 to 4 (mean of sum per dark phase, n = 4 dark phases). **c** Average number of sniffing events with familiar mice during the dark cycles of days 1 – 4 (2-tailed unpaired Mann-Whitney test: U = 1, p = 0.0571, n = 4 dark cycles). **d** Average duration of sniffing events with familiar mice during dark cycles of day 1 to day 4 (2-tailed unpaired t-test: t(6) = 8.544, p = 0.0001, n = 4 dark cycles). **e** Visual representation of the test paradigm on days 4 to 6 of behavioral experiments. **f** Social Score of individuals during 6-8pm on days 4 to 6 of social arena experiments (2-tailed mixed-effects one-way ANOVA with Geisser-Greenhouse correction and Fisher’s LSD posthoc test, F(2.007, 14.85) = 4.554, p = 0.0286). **g** Locomotion Score of individuals during 6-8pm on days 4 to 6 of social arena experiments (2-tailed mixed-effects one-way ANOVA with Geisser- Greenhouse correction and Fisher’s LSD posthoc test, F(2.595, 19.21) = 12.43, p = 0.0002). **h-j** Difference Scores (DS) of social sniffing during the initial 2h of dark phase, calculated in 5min bins. **h**) DS for interaction with mixed (day 6) vs familiar (day 5) cohort (2-tailed Mann Whitney t-test, U = 149, p = 0.0036). **i**) DS for interaction with familiar (day4) vs mixed cohort (day5). **j**) *left* DS for interaction with familiar mice before (day4) and after (day6) 24h separation (2-tailed unpaired t-test, t(46) = 3.219, p = 0.0024). *right* Line graph shows the mean DS during initial 30min of social interaction period (2-tailed RM two-way ANOVA with Geisser- Greenhouse correction and Fisher’s LSD posthoc test, time x group F(5, 75) = 1.482, p = 0.2059). Data are males, Vh: n = 8 mice and hM4Di^+^: n = 9 mice. Bar graphs show mean ± SEM. ns = not significant, * p < 0.05, ** p < 0.01, *** p < 0.001. Statistical details are shown for significant changes only; all details are provided in statistical details file.

In this study, we demonstrated that CA2 inhibition induces long-lasting disruptions in neuronal activity and connectivity of frontal, thalamic and hippocampal brain circuits with consequences for social behavior. fUS imaging demonstrated brain-wide acute hypoactivity and lasting functional hypoconnectivity of the frontal network due to CA2 inhibition. In particular, the hippocampal-fronto-thalamic connectivity appeared to depend on intact excitatory CA2 activity. Mice showed subtle impairments in their social interaction with familiar mice persisting for hours.

We provided the first *in vivo* proof of dCA2’s regulatory power over the activity of the dorsal and ventral hippocampal subfields^15,16^, although more drastic changes on the connectivity level were expected given CA2’s high degree of intrahippocampal connectivity^9,15,16^. Moreover, we support earlier reports on a functional dCA2-vCA1-mPFC axis^4,17,18^, previously attributed to social memory ^4,17,18^. Surprisingly, thalamic activity was strongly reduced upon CA2 inhibition as well as the mPFC-thalamus FC. As synaptic connectivity between dCA2 and thalamus has not been described yet^9^, we propose a dCA2>vCA1>mPFC>thalamus cascade, considering previously reported dCA2>vCA1>mPFC^4,11,17^ and mPFC-thalamus circuits^12^.

Social cognition has recently been mapped to a frontothalamic circuit^12^ , thus reduced mPFC-thalamic FC may contribute to changes in social interaction and memory, described here and in previous studies.

Considering the established role of CA2 in social recognition memory^3,4^ , our data might represent a social memory impairment in hM4Di^+^ mice while vehicle control mice show intact memory – potentially manifested due to alteration in the above-mentioned brain circuits. The repetitive “sniff and hop”-like short-term interaction of hM4Di^+^ mice with conspecifics may also represent a subtle form of social memory impairment. It remains unclear whether the 24 h separation or the encounter with novel animals exacerbated interaction difficulties. CA2 has been associated with aberrant salience^19^, a maladaptive behavior of shifting attention, in patients with schizophrenia. Hence, our observation might be alternatively explained as an aberrant shift in attention towards conspecifics resulting in repetitive sniffing.

CA2 inhibition recapitulates several network-level endophenotypes characteristic of psychiatric disorders. Specifically, disrupted hippocampal–medial prefrontal cortex (HPC– mPFC) and mPFC–thalamic connectivity, both consistently implicated in schizophrenia and related disorders^20,21^, emerged following CA2 silencing. In addition, CA2 inhibition induced a hypoconnected frontal network state resembling alterations observed in chronic schizophrenia patients^22^ and preclinical models of subchronic NMDAR hypofunction^23^. These disruptions are linked to key disease-relevant endophenotypes, including deficits in social cognition, impaired executive control, dysfunctional thalamo-cortical processing, and abnormal large-scale network coordination^1,10,21^.

Limitations of this study are the sedation during fUS acquisitions, potentially affecting neuronal activity and FC^24^, and limits in spatiotemporal resolution of the imaging technique, which may explain the lack of identified changes in hippocampal microcircuits^15,16^. Despite the advantages of the social arena^14^ , the comparability of data with other studies is limited. And finally, an antipsychotic action by CNOs metabolite clozapine needs to be considered^25^.

Collectively, our findings identify CA2 integrity as a critical determinant of brain-wide network stability relevant to psychiatric disease mechanisms.

## Declaration of competing interest

The authors declare no conflicts of interest.

Results of this work were previously published as part of a dissertation (https://doi.org/10.18725/OPARU-59196).

## Author contribution

Alessa A. Franz: Conceptualization, Investigation, Formal analysis, Writing – original draft, Writing – review & editing. Tudor M. Ionescu: Formal Analysis, Writing – review & editing. Dennis Kätzel: Conceptualization, Supervision, Writing – review & editing. Bastian Hengerer: Conceptualization, Supervision, Writing – review & editing.

## Acknowledgments

The authors thank Dr. Benjamin Strobel and his laboratory for providing the AAVs used in this study; Aileen Reich, Nancy Kötteritzsch and the Biological Laboratory Service, Boehringer Ingelheim for their support with animal care taking; Olena Sakk from the Transgenic Core Facility of Ulm University for cryopreservation and rederivation of the experimental animals; the Tierforschungszentrum of Ulm University for their support regarding the mouse transfer; Dr. Romina Schnegotzki, Dr. Klaus Brilisauer and their teams for the bioanalysis of plasma samples, Shania Smith for her support and Marti Ritter for bioinformatic input.

## Appendix A

Supplement, Statistical details

## Methods

### Animals

All experiments were authorized by the governmental animal ethics committee (Regierungspräsidium Baden-Württemberg, Tübingen, Germany; license number 20-034-G) and performed in compliance with local animal care guidelines and with the Association for Assessment and Accreditation of Laboratory Animal Care (AAALAC) regulations.

The B6.Cg-Tg(Amigo2-cre)1Sieg/J BAC-transgenic mouse line was originally obtained from The Jackson Laboratory (MA, US; stock number 030215) and bred in-house at Ulm University. Male and female experimental animals were obtained by rederivation from frozen embryos in the transgenic core facility of Ulm University and genotyped for hemizygous Cre (Amigo2- Cre^+^)^3^. Mice were group-housed (2 - 4 animals per cage) in Makrolon Type III cages containing standard enrichment (one house, one tunnel, a piece of wood and nesting material) in a temperature (22 °C ± 2 °C) and humidity (± 45-60%) controlled room and subjected to a 12 h light/dark cycle (light cycle starting at 6am). Food and water were supplied *ad libitum*.

### Virus constructs

Specific targeting of CA2 pyramidal neurons in Amigo2-Cre^+^ mice was achieved by bilateral infusions of AAV8 vectors at final titers 2.07 *10^12^ VG/ml (diluted in 1x PBS) below and above the pyramidal cell layer of dorsal CA2 (dCA2; see stereotaxic surgeries). The viral constructs contained a double-floxed inverted open reading frame (DIO) design for Cre-dependent expression of the DREADD hM4Di fused to the fluorescent reporter mCherry (hM4Di^+^ group; AAV8-hSyn-DIO-hM4Di-mCherry-WPRE) or mCherry alone (AAV8-hSyn-DIO-mCherry-WPRE). Vectors were obtained from in-house production.

### Stereotaxic surgeries

Stereotaxic virus injections for behavioral studies were performed on 9- to 11-week old mice and for fUS imaging on 7- and 10-week-old mice. After pre-surgical treatment with Metacam (Meloxicam 5 mg/kg, Boehringer Ingelheim Vetmedica GmbH), mice were deeply anaesthetized (isoflurane, max. 1.5%, Virbac Tierarzneimittel GmbH), placed in a stereotaxic frame (model 1900, KOPF Instruments) and local anesthesia (Carbostesin® 0.25%, Aspen) applied prior to opening the skull. AAV8-mCherry or AAV8-hM4Di-mCherry (see virus constructs above) were delivered bilaterally to dCA2 via a microliter syringe (model 62 RN, Hamilton) at the following coordinates: AP -2.1 mm, ML ±2.4 mm, DV -2.3 mm (volume: 600 nl) and -2.1 mm (volume: 400 nL) from skull surface at Bregma (defined as triangular midpoint above Bregma). Infusion speed was 100 nl/min with a post-injection waiting period of 5 min per site and additional 5 min waiting 0.05 mm dorsal to the last DV-injection. Surgical procedures were performed in compliance with German standards and the animal well-being during and after surgery guaranteed.

Mice received clozapine-N-oxide (CNO; CNOCl2, in-house production; 2 mg/kg) dissolved in drinking water starting from 2 weeks post-surgery. Water bottles were protected from light via aluminum foil and exchanged with freshly dissolved compound every second day.

### Behavioral assessment

Social behavior of male Amigo2-Cre^+^ (for Vh, n = 8 mice; for hM4Di^+^, n = 9 mice) mice was assessed in the social arena apparatus as previously described^14^ 3 weeks post-injection. We equipped the arena with a big nest containing nesting material and two small nests, with and without nesting material (see Fig. S6h_i_). Mice were provided with CNO in drinking water (see Stereotaxic surgeries) and food *ad libitum*. All arenas were equally equipped. Experiments were conducted on a 12h light/dark cycle, with the dark phase starting at 6pm, for an experimental period of 9 consecutive days. Homogenous groups of maximum 4 animals per arena were formed and maintained throughout (with exception, see social test paradigm). Animals were habituated to the arena for three consecutive days (Hab day 1 to Hab day 3), followed by 3 days of social interaction (day 1 to day 3) and 3 days of social test paradigm (day 4 to day 6). Habituation on Hab day 1 started at 4pm, otherwise start of 22h video recordings were at 6pm. Mice stayed in the arena during the whole experimental period and were only moved back to the home cage (together with area mates) during an inter-trial interval (ITI) of max. 2h for regular arena cleaning.

The social interaction was scored by the time mice spent with social sniffing at head, body and genitals of the interaction partner or social follow. Social scores were calculated, consisting of social sniffing and social follow. Locomotion was quantified as time mice spent in locomotion in the periphery and center of arena. Locomotion scores were calculated from the duration of locomotion (periphery and center) and time mice spent in the arena.

### Social test paradigm

After 4 days of interaction in homogenous familiar groups, heterogenous same-sex groups of familiar and novel mice were generated and maintained in the arena for 22h on day 5 of experiment, starting at 6pm. Each arena housed hM4Di^+^ and control animals, both encountering a familiar mouse and two novel mice (see Fig. 3b). On day 6, mice were placed back into familiar groups. Each arena including nests was cleaned and disinfected during a 2h ITI between sessions, lasting from 4pm to 6pm, to avoid an influence of odor on the experimental outcome. Social groups were maintained during the ITI and transferred into the home cage. Social interaction and locomotion were scored as described above. Difference Score calculations from Hitti and Siegelbaum^3^ were adapted to fit our behavioral assessment, thus scores were calculated for the time spent in social interaction during the initial 2h of interaction between distinct days of the social test paradigm.

### Automatic behavior analysis

Video material and RFID-information from the behavioral tracking were analyzed as previously described ^14^. In brief, the RFID-Assisted Social Scan Software package was used to create an arena layout defining various regions (like center and periphery, see Fig. S6h_i_) within the arena, in which the behavior of mice was analyzed. Pre-defined parameters describing social and non-social behavior events were fed into the software. Videos were automatically scanned for those behaviors in batch mode. The data were exported in csv format and processed by MATLAB (MathWorks, version EU 2022b) and Prism 9 (GraphPad Software, version 9).

### Functional Ultrasound Imaging

Cerebral blood volume (CBV) of mCherry and hM4Di-mCherry injected Amigo2-Cre^+^ male (n = 11; for Vh n = 5 mice; for hM4Di n = 6 mice) and female (n = 13; for Vh n = 7 mice; for hM4Di n = 6 mice) mice were measured by functional ultrasound imaging (ICONEUS One, ICONEUS, France). Starting 3 weeks after viral injection, mice were scanned once per week for 90min over a time course of 3 consecutive weeks. During the initial scan, an acute CNO injection (5 mg/kg, i.p.) was applied to mice after 40 min of baseline recording. From the following day on, mice underwent chronic CNO treatment via drinking water (as described above) until end of experiment. fUS scans under chronic CNO influence were performed on days 6 and 13 of chronic treatment.

Mice were scanned using the IcoPrime – 4D MultiArray Probe^8^ (ICONEUS, France) under slight sedation (0.09mg/kg subcutaneous dexmedetomidine hydrochloride bolus (Sigma) + 3% isoflurane in oxygen prior to the scan and 0.5% isoflurane + slow dexmedetomidine infusion (0.06 mg/kg/h) during scan), head-fixation (Stereotaxic frame model 1900, KOPF instruments) and stable body temperature (35 - 36.9 °C) conditions. The fUS probe was placed ∼1 mm above the shaved scalp, which was covered with air bubble free ultrasonic gel (Parker Laboratories, Inc., #01-08). Bilateral, coronal scans spanning from frontal cortex towards midbrain were acquired with a frequency of 15 MHz (acquisition of 4 x 200 compounded frames at a frame rate of 500 Hz in 0.4 s per probe position; one 4 x 4 slice frame = 2.4 s). An angiographic image, acquired for each session and animal, was co-registered to the Allen brain atlas integrated within the IcoScan Software (ICONEUS, France). Data were acquired starting after a minimum of 10 min after the onset of 0.5% isoflurane administration. After scanning, mice received subcutaneous atipamezole (Alzane, Zoetis) injection at a dose of [administered dexmedetomidine infusion dose + 90]/2 to antagonize the sedation and recovery was monitored.

A few days after the final fUS scan, blood samples were taken from the heart immediately before perfusion (according to protocol above), and plasma concentration of the CNO’s metabolite clozapine^25^ was analyzed in-house.

### Data preprocessing and analysis

Acquired fUS images were saved in Nifti format and underwent the following preprocessing steps using SPM12 software (<u>SPM12 Software - Statistical Parametric Mapping</u>): realignment, co-registration to the Allen mouse brain atlas, spatial smoothing using a 0.2×0.2×0.2 mm^3^ Gaussian kernel and baseline linear detrending for CBV calculations. Regional time courses were extracted from the preprocessed images.

The delineation of all analysed regions is based on the Allen Brain atlas. Thalamic nuclei were grouped in the following 5 subgroups: ilmed (midline group, medial group and intralaminar nuclei of dorsal thalamus), anterior, lateral and ventral thalamic nuclei as well as MD. Details are provided in the Source data file. Classification of the thalamic regions is visualised by using the web-based atlas provided at Kimab Unified Anatomical Atlas (https://kimlab.io/brain-map/atlas/atlas_viewer.html?s=60_AP-1.6).

#### Regional activity

Using the preprocessed images, acute chemogenetically induced CBV alterations were analyzed on two distinct spatial levels. For investigating the CBV changes at regional level, images were normalized to the baseline period defined as the final 10 minutes before CNO application (minute -10 to 0 of acute CNO scan). To generate relative CBV (rCBV) time courses, each regional time course was normalized to its respective average value over this time period.

To investigate CBV alterations at voxel level, a first-level general linear model (GLM) analysis was applied to the individual scans using the pseudo block approach reported for phMRI using SPM 12^26^. The following blocks were used: minute -5 - 0 as baseline block and 0 - 10 min, 10 – 20 min, 20 – 30 min, 30 – 40 min and 40 – 50 min as post-CNO blocks. Following GLM parameter estimation, the data were interrogated using contrast vectors between each post-CNO application block, on one side, and the baseline block on the other side, in order to generate statistical parametric maps. These subject-level maps were then used to generate group-level effects using a second-level analysis.

#### Functional connectivity

Prior to functional connectivity (FC) computation, all regional time courses were bandpass-filtered (0.01-0.2 Hz). Using these filtered time courses, Pearson’s correlation coefficients were computed for all pairs of regional time courses to generated ROI-to-ROI FC matrices.

Group-level FC matrices were generated by transforming all single-subject Pearson’s r correlation coefficients to Z-scores using Fisher’s Z-transformation and averaging those to group-level. The period minute -35 to 0 of the acute CNO scan were defined as the baseline and used for computation of both acute and chronic CNO effects. For the acute CNO acquisition, an ANCOVA between control and hM4Di^+^ group was performed, using the respective baseline values as covariates. For all chronic CNO measurements, a linear mixed model was used to compare minutes 5 to 40 of both chronic CNO scans with the baseline for both groups.

Exclusion criteria were extreme changes in body temperature during a scan, reaching the animal welfare endpoint criteria of the experiment and unstable baseline recording. The data were processed and plotted by MATLAB (MathWorks, version EU 2022b), Prism 9 (GraphPad Software, version 9) and OriginPro9 (OriginLab, version Pro9).

### Perfusion and Immunohistochemistry

Histology was performed as previously described by us^23^. Mice were deeply anesthetized with pentobarbital and transcardially perfused with 1x PBS (Gibco) followed by 4% paraformaldehyde (PFA; Electron Microscopy Sciences) in 1x PBS. Brains were post-fixed in 4% PFA overnight at 4 °C and afterwards transferred to 30% sucrose. Using a freezing stage sliding microtome (Epredia™ HM450), 30µm coronal sections were prepared. Free floating histology was performed on two consecutive sections (at -1.9 mm from bregma) per animal covering the dorsal hippocampus. Slices were blocked with 0.3% Triton X-100 (Sigma), 10% normal goat serum (Cedarlane) and 1% BSA (Dianova; IgG and protease free) in 1x PBS for 2 h at room temperature under steady agitation. Primary antibodies were diluted in carrier solution containing 0.3% Triton X-100, 1% normal goat serum and 1% BSA and sections incubated overnight at 4 °C under gentle agitation: guinea pig anti-Parvalbumin (Synaptic Systems, #195004, 1:500), rabbit anti-PCP4 (Sigma-Aldrich, #HPA005792, 1:200). Afterwards, sections were washed for about 2 h with periodic exchange of washing solution (0.3% Triton X-100, 1% normal goat serum and 1%BSA (Sigma) in 1x PBS). The sections were then incubated for 2h (room temperature and under gentle agitation) with the following secondary antibodies diluted in carrier solution: goat anti-rabbit Alexa Fluor 488 (Invitrogen, #A11070, 1:1000), goat anti-guinea pig Alexa Fluor 647 (ThermoFisher, #A21450, 1:1000). After washing five times for 15 min, sections were mounted in 24-well glass bottom plates (Sensoplate, Greiner), air dried and imbedded with aqueous mounting medium containing DAPI (Fluoroshield™ with DAPI, Sigma).

### High-throughput confocal microscopy

Fluorescent images were obtained with an Opera Phenix (PerkinElmer) high-throughput microscope using the 63x objective in confocal mode. For each section, ∼480 visuals fields (200 µm x 200 µm individual field size) covering the dorsal hippocampus were obtained comprising a stack of 6 planes with a distance between single planes of 1 µm. For optimal image quality, a laser power of 100% and laser exposure times attuned for every antibody were used.

### Statistical analysis

Statistical data analysis was performed in Prism9 (GraphPad Software, version 9). Normal distribution of data was tested using the Shapiro-Wilk test. Statistical tests used for each data set are presented in the figure legend. In general, in-group comparison between days was performed via repeated measure ANOVA. Comparisons between test groups were performed via t-test. Statistical tests for the fUS data set are described in the data analysis.

---

**Fig. S1.**
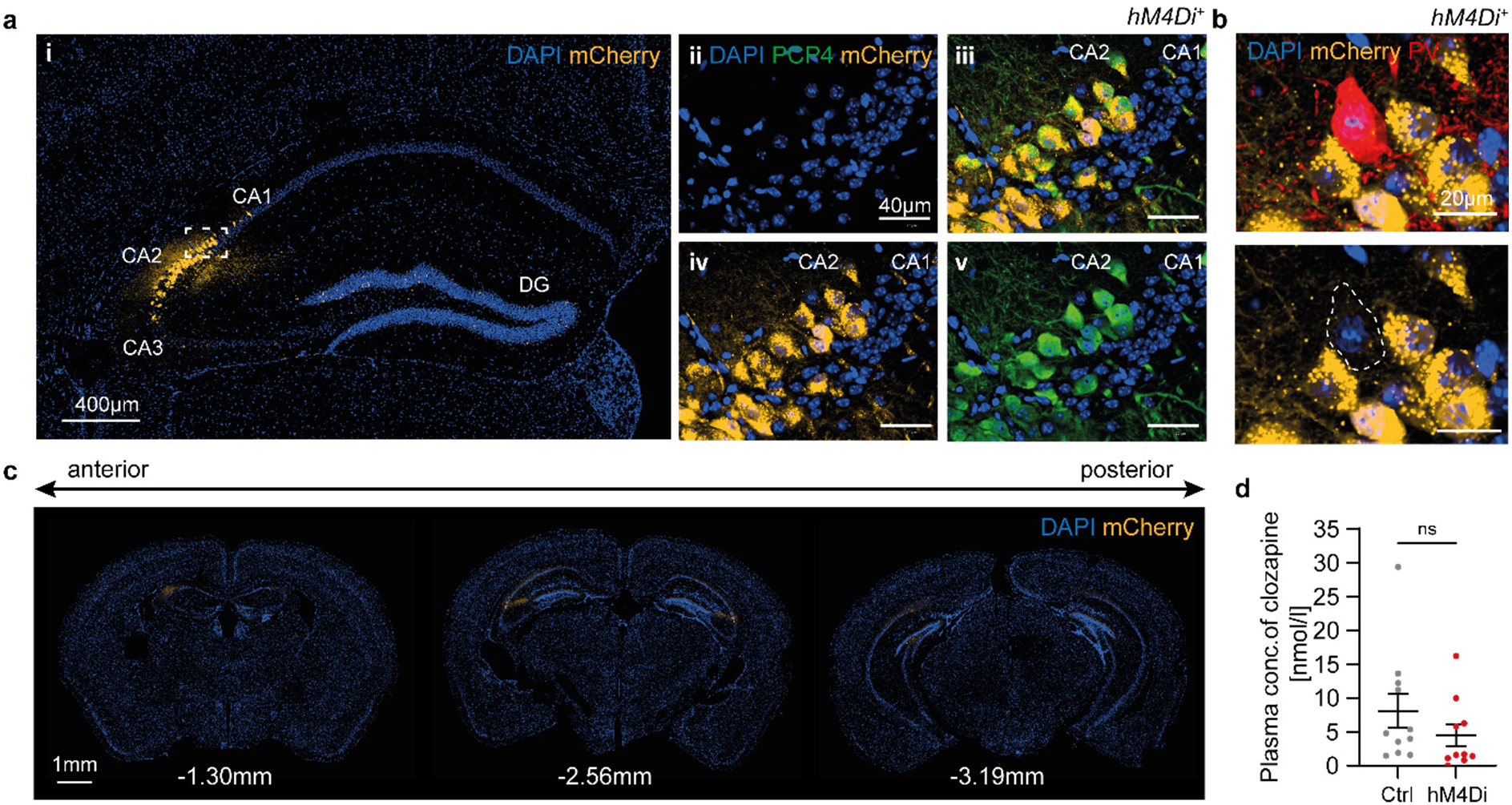
Specific hM4Di^+^ expression in CA2 pyramidal neurons for acute and chronic CA2 inhibition. **a** Representative confocal image of hM4Di-mCherry (orange) infused dorsal hippocampus (**a_i_**). Scale bar = 400 µm. Zoom in (**a**_ii-v_) of dashed area in (**a_i_**) shows the expression of mCherry and PCP4 (green) in the CA2 subregion. Scale bar = 40 µm. **b** Expression of hM4Di-mCherry and inhibitory parvalbumin-interneuron in dorsal hippocampus. Scale bar = 20 µm. **c** Viral spread across dorsal towards ventral hippocampus. Coordinates of slices are indicated by distance from Bregma in mm. Scale bar = 1mm. DAPI (blue) indicated cell nuclei. **d** Clozapine concentration in blood plasma of mice after exposure to CNO for 3 weeks. Data point represents individual values (for Vh, mean concentration: 8.0888 ± 2.508 nmol/l, n = 11 mice; for hM4Di^+^, mean concentration: 4.483 ± 1.644 nmol/l, n = 10 mice; 2-tailed unpaired Mann-Whitney t-test: U = 34, p = 0.1517).

**Fig. S2.**
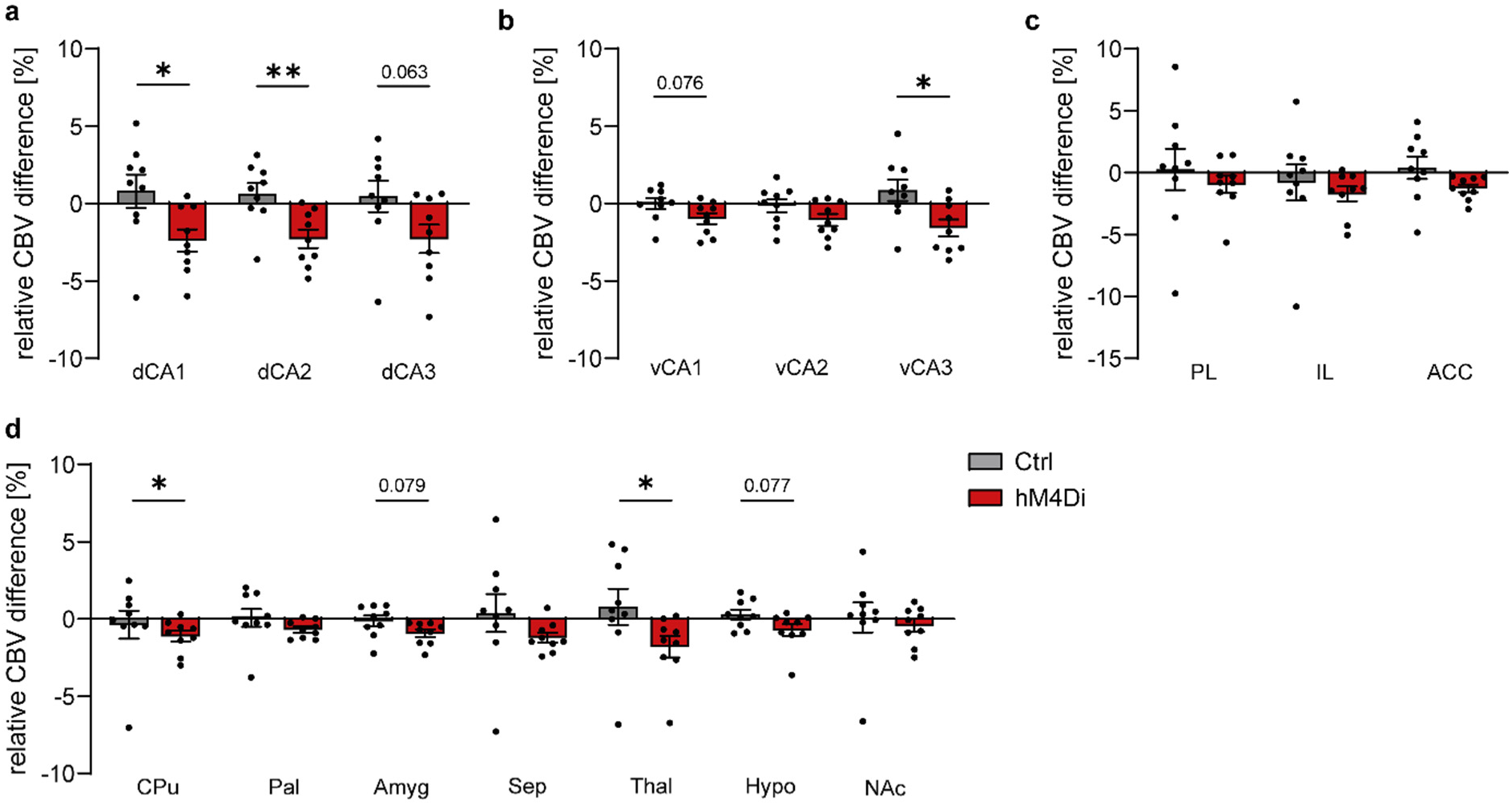
Effects of acute CA2 inhibition on the regional activity of distinct brain regions. Scatter plots show the percental change in cerebral blood volume (CBV) during the last 10 min of acute CNO fUS acquisition in **a** dorsal hippocampus (2-tailed unpaired t-test: dCA1 t(16) = 2.476, p = 0.0248; dCA2 t(16) = 3.286, p = 0.0047), **b** ventral hippocampus (2-tailed unpaired t-test: vCA3 t(16) = 2.755, p = 0.0141), **c** prefrontal areas and **d** further brain regions (2-tailed unpaired Mann-Whitney t-test: CPu U = 16, p = 0.0315; Thal U = 15, p = 0.0244). **a-c** are the raw data of Fig. 1c. Bar graphs show mean ± SEM. Each data point represents a measured animal. Only significant changes are shown, *p < 0.05, ** p < 0.01. All statistical details are provided in statistical details file.

**Fig. S3.**
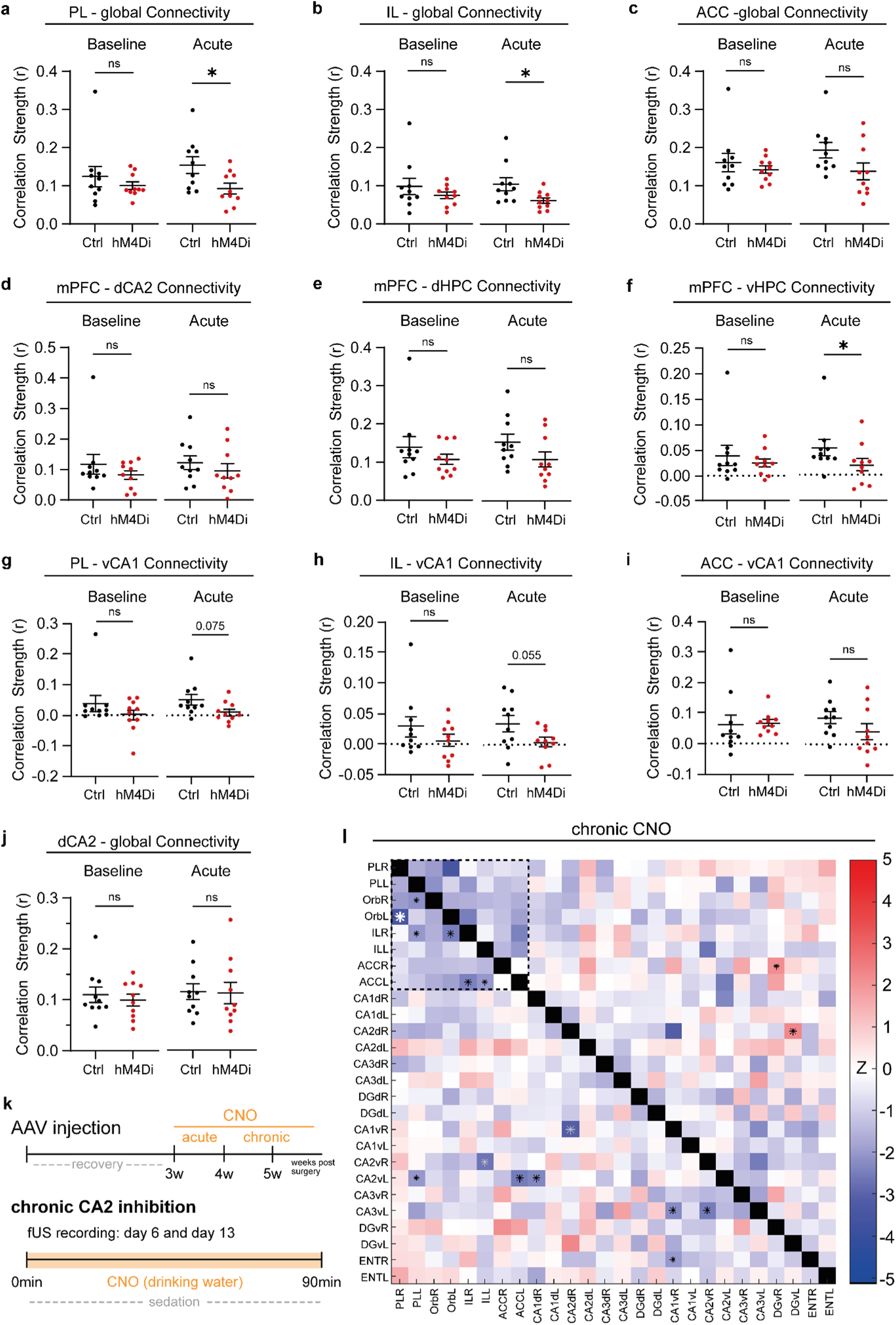
Functional HPC-mPFC connectivity after acute and chronic CA2 inhibition. **a-c** Global functional connectivity strength as average Pearson’s r correlation during baseline and acute CNO of (**a**) PL (acute, 2-tailed unpaired t-test: t(18) = 2.378, p = 0.0287), (**b**) IL (acute, 2-tailed unpaired Mann-Whitney t-test: U = 19.5, p = 0.0196) and (**c**) ACC. **d-f** Functional connectivity strength between the mPFC and (**d**) dCA2, (**e**) dorsal hippocampus or (**f**) ventral hippocampus (2-tailed unpaired Mann-Whitney t-test: U = 22, p = 0.0355). **g-i** Functional connectivity strength between vCA1 and the prefrontal areas (**g**) PL, (**h**) IL (2-tailed unpaired t-test: t(18) = 2.052, p = 0.0550), (**i**) ACC. **j** Global functional connectivity strength of dCA2. **k** Timeline of entire fUS experiments in the upper panel and experimental overview of the chronic CNO recordings in the lower panel. **I** Heatmap of functional connectivity changes between regions of the frontal cortex and hippocampus in hM4Di^+^ mice relative to controls. Dashed square indicates the mPFC-cluster of hypoconnectivity. Statistical difference is shown via Z-scores. Z < 0 is control > hM4Di^+^ (blue color), Z > 0 is Control < hM4Di^+^ (red color). *black: p < 0.05, *grey: p < 0.01, *white: p < 0.001. Data are males, n = 10 mice per group. Each data point represents a measured animal. Error bars are SEM. *p < 0.05, ** p < 0.01. Statistical details are shown for significant changes only; all details are provided in statistical details file.

**Fig. S4.**
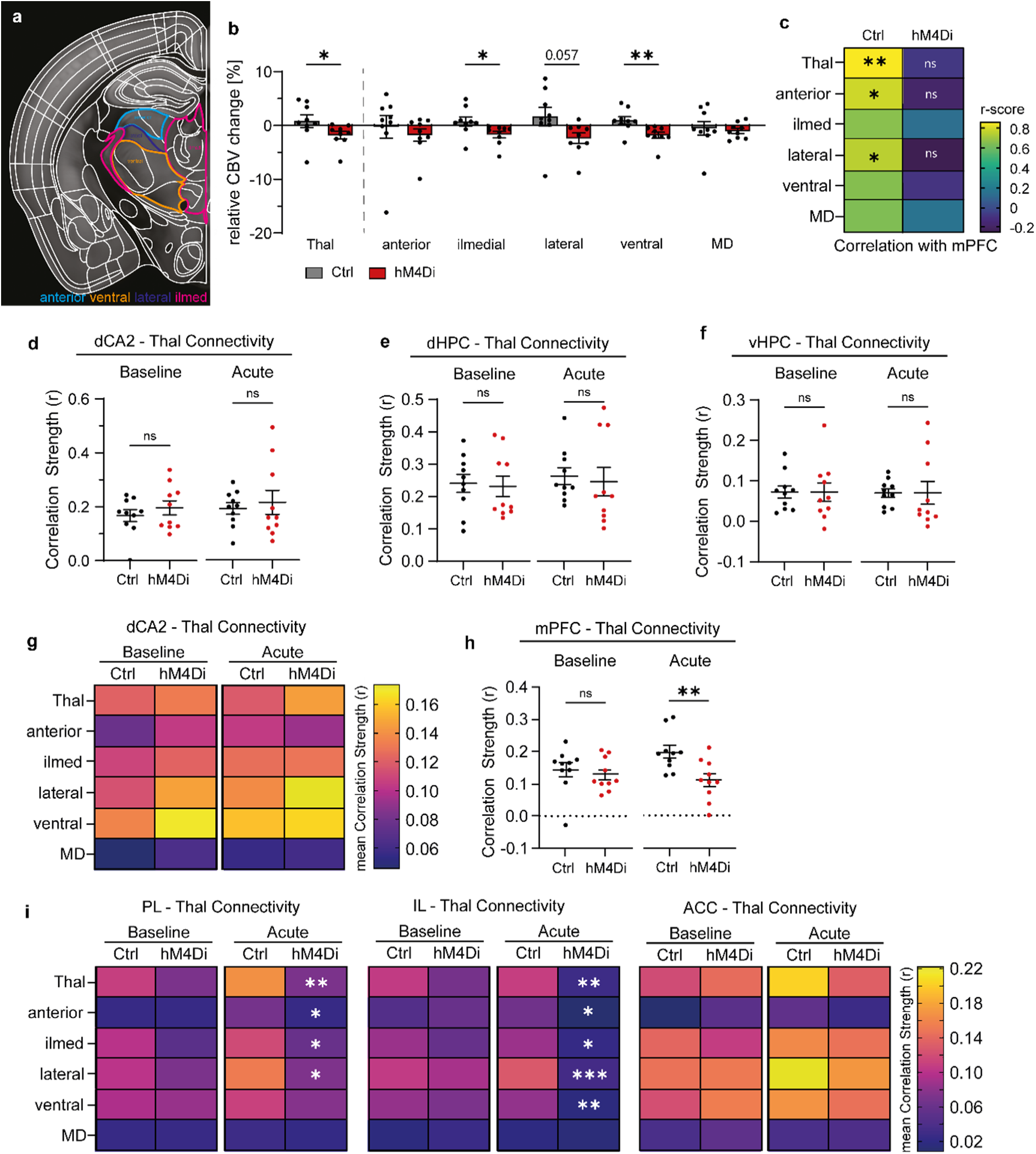
Functional thalamic connectivity after acute CA2 inhibition. **a** Exemplary grouping and delineation of thalamic nuclei. Image adopted from <u>Kimab Unified</u> <u>Anatomical Atlas</u>; Groups are classified according to the Allen Brain Atlas, see methods. Pink = ilmed group (midline group, medial group and intralaminar nuclei of dorsal thalamus); light blue = anterior group; dark blue = lateral group; orange = ventral group; MD not shown. **b** Percental change in CBV of the entire thalamus and thalamic nuclei during the last 10 min of acute CNO scan (2-tailed unpaired Mann-Whitney t-test: Thal U = 15, p = 0.0244; ilmedThal U = 12, p = 0.0106; vThal U = 8, p = 0.0028). **c** Color-coded correlation in rCBV between mPFC and thalamic nuclei for Vh and hM4Di^+^ mice following acute CNO (minute 40-50; Pearson r correlation and 2-tailed unpaired t-test for Vh, Thal: r = 0.8571, p = 0.0032, aThal: r = 0.8571, p = 0.0032, lThal: r = 0.7458, p = 0.0210; for hM4Di^+^, Thal: r = -0.0844, p = 0.8291, aThal: r = -0.1853, p = 0.6332, lThal: r = -0.2572, p = 0.5041). **d-f** Functional connectivity of (**d**) dCA2, (**e**) dHPC and (**f**) vHPC with the entire thalamus during baseline and acute CNO. **g** Heatmap of functional connectivity between dCA2 and thalamic nuclei as r-score for 40-50 minutes after acute CNO. **h** Functional connectivity of mPFC with the entire thalamus (acute, 2-tailed unpaired t-test: t(18) = 3.773, p = 0.0014). **i** Heatmap of functional connectivity between thalamic nuclei and the mPFC subregions PL (*left*), IL (*middle*) and ACC (*right*) before and after acute CNO (acute, 2-tailed unpaired t-test: for PL-Thal t(18) = 3.657, p = 0.0018, for PL-ilmed t(18) = 2.102, p = 0.0499, for PL-lateral t(18) = 2.493, p = 0.0226, for IL-Thal t(18) = 2.961, p = 0.0084, for IL-lateral t(18) = 3.965, p = 0.0009, IL-ilmed t(18) = 2.645, p = 0.0164, IL-ventral t(18) = 2.944, p = 0.0087; 2-tailed Mann-Whitney t-test: for PL-anterior U = 18, p = 0.0147, for IL-anterior U = 17, p = 0.0115). Data points are individual mice. n = 10 mice per group for FC and n = 9 mice per group for rCBV. Error bars are SEM. *p < 0.05, ** p < 0.01, *** p < 0.001. Statistical details are shown for significant changes only; all details are provided in statistical details file.

**Fig. S5.**
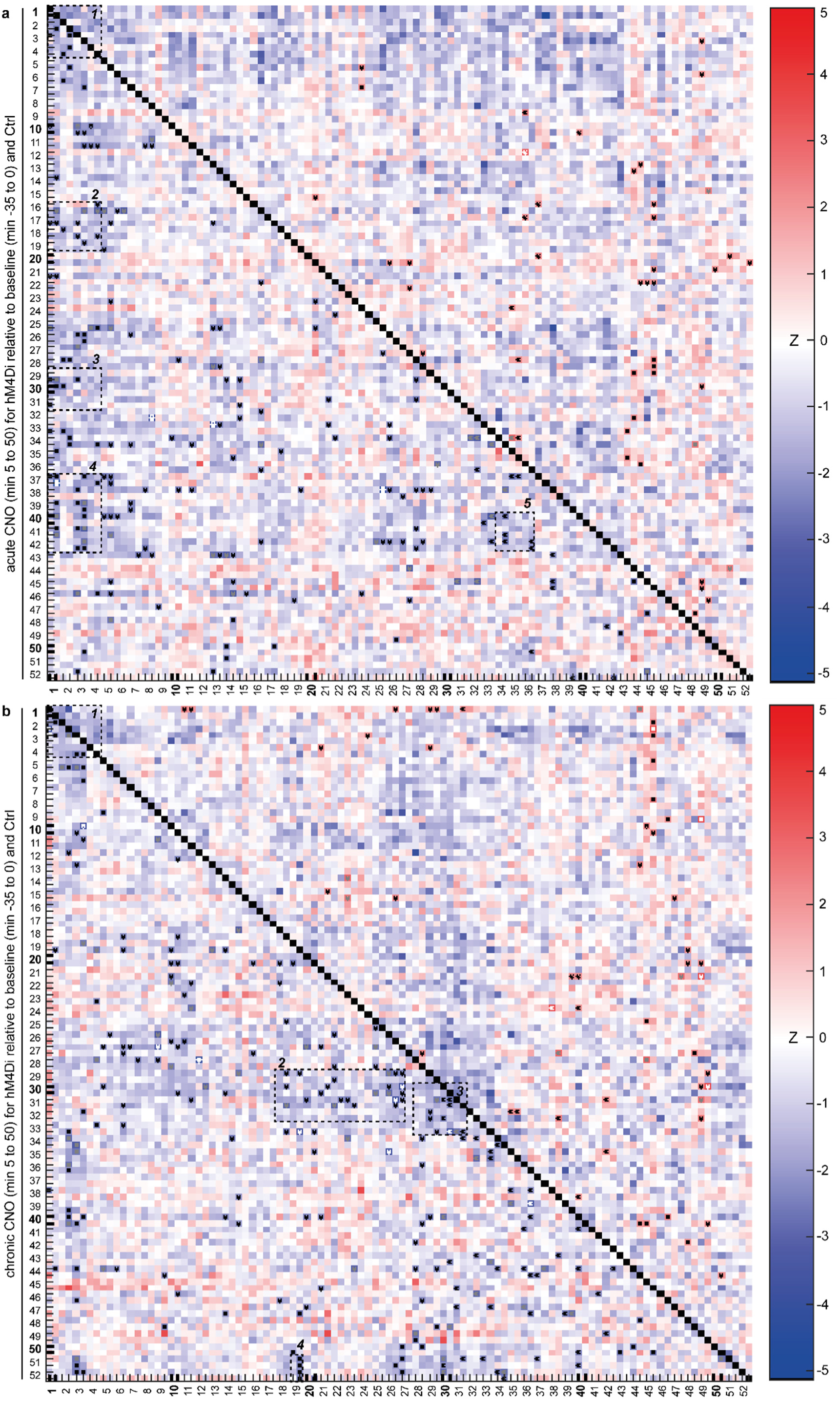
Functional connectivity heatmaps following acute and chronic CA2 inhibition. **a** Effects of acute CNO application (minutes 55 to 90) on the functional connectivity between all measured brain regions in hM4Di^+^ mice relative to controls. Clusters of significantly reduced connectivity between brain regions are indicated by numbered dashed squares: 1: frontal cortex; 2: HPC – frontal cortex; 3: amygdala – frontal cortex; 4: thalamus – frontal cortex; 5: septum – thalamus. **b** Effects of chronic CNO application (minutes 5 to 40) on the functional connectivity between all measured brain regions in hM4Di^+^ mice relative to controls. Dashed squares indicate clusters of significantly reduced connectivity: 1: frontal cortex; 2: amygdala – HPC; 3: Amygdala-Amygdala-Pal-NAc; 4: vCA2 – SC/IL. Statistical difference is shown via Z-scores. Z < 0 is control > hM4Di^+^ (blue color), Z > 0 is Control < hM4Di^+^ (red color). *black: p < 0.05, *grey: p < 0.01, *white: p < 0.001. Otherwise, ns = not significant, *p < 0.05, ** p < 0.01. Regions are: 1: PL, 2: Orb, 3: IL, 4: ACC, 5: M2, 6: M1, 7: SS, 8: VIS, 9: AUD, 10: OLF, 11: PIR, 12: TEa, 13: RSP, 14: CA1d, 15: CA2d, 16: CA3d, 17: DGd, 18: CA1v, 19: CA2v, 20: CA3v, 21: DGv, 22: ENT, 23: ECT, 24: PERI, 25: SUB, 26: CPu, 27: Pal, 28: NAc, 29: Amyg, 30: BLA, 31: CEA, 32: Ins, 33: CLA, 34: Sep, 35: MS, 36: LS, 37: Thal, 38: MD, 39: aThal, 40: ilmedThal, 41: lThal, 42: vThal, 43: Hypo, 44: SUM, 45: PVH, 46: LHA, 47: mHypo, 48: SN, 49: VTA, 50: PAG, 51: SC, 52: IC. Detailed directory of x- and y-labels is provided in statistical details file.

**Fig. S6.**
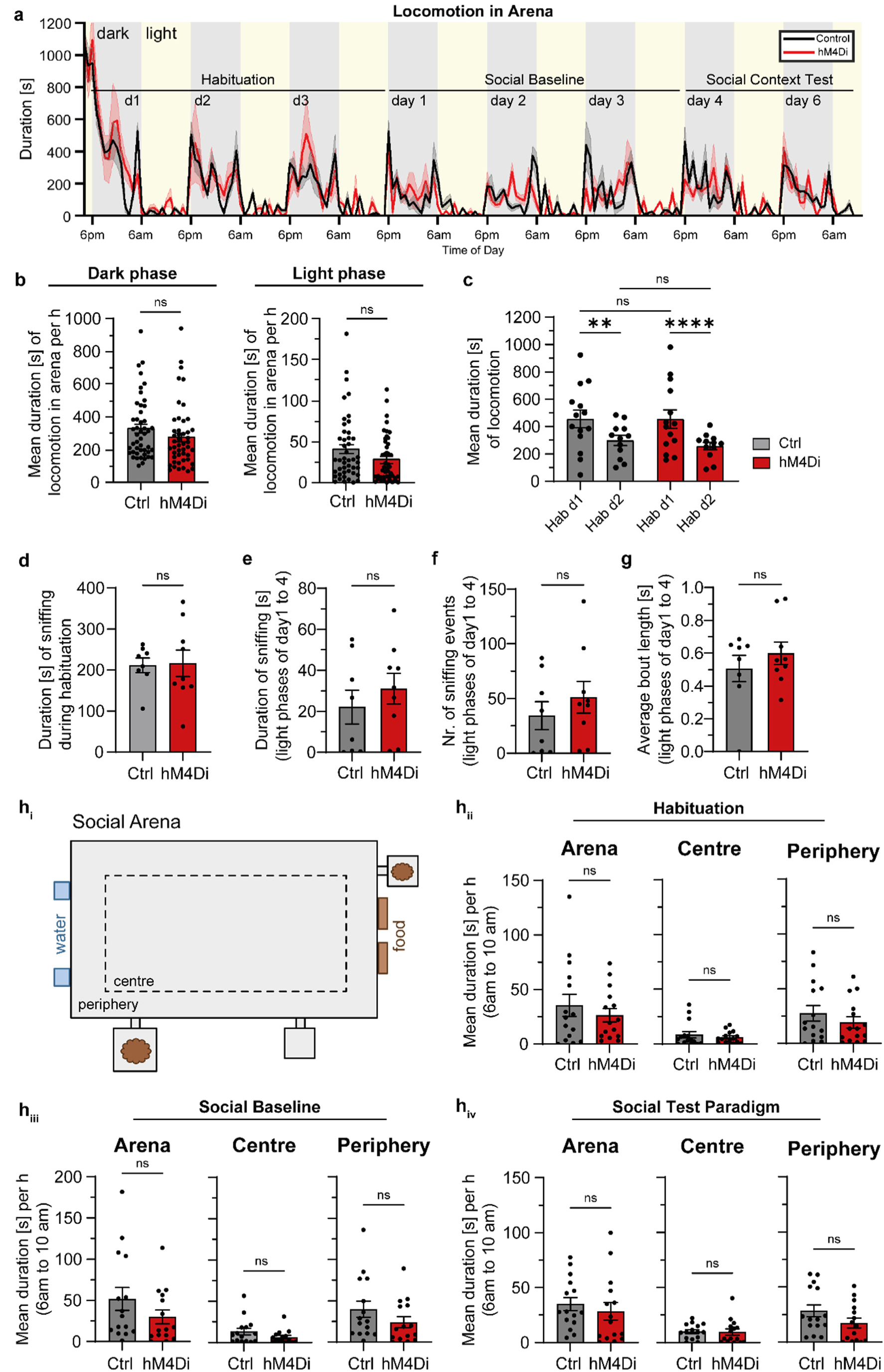
Habituation, non-social and anxiety-related behavior of mice in social arena. **a** Representative line graph shows locomotion duration in arena of control and hM4Di^+^ males in familiar groups during the time course of experiment (males, Ctrl: n = 4, hM4Di: n = 4; familiar groups, 4 mice per arena; mean duration per h ± SEM). **b** Average time spent with locomotion per hour during initial 5h of dark phase (*left*) and light phase (*right*) on all experimental days (n = 47h, time period is 6pm to 11pm, exception: Habituation day1 = 4pm to 11pm (see methods), data point is group mean per hour, 2-tailed unpaired Mann-Whitney test). **c** Mean duration of locomotion during two consecutive days of habituation (data points are group mean per hour, 2-tailed 2-way ANOVA with Fisher’s LSD posthoc test: F (3, 425) = 10.14, p < 0.0001). **d** Time spent with social sniffing during the nocturnal habituation period (data point is sum per individual, 2-tailed unpaired t-test). **e** Average of accumulated time mice spent sniffing with familiar mice light phase (6am-4pm) of days 1 to 4 **f** Average number of sniffing events with familiar mice during the light phase of days 1 to 4 **g** Average duration of sniffing events with familiar mice during light phase of day 1 to day 4. **h** Average duration of locomotion per hour in centre, periphery and total arena during 6-10 am of distinct experimental phases. **h_i_** Compartmentalisation of centre and periphery in arena. **h_ii_** habituation (Hab days 1-3) **h_iii_** social baseline (days 1-3) and **h_iv_** social interaction paradigm (days 4-6). Males, n = 8 mice for Ctrl and n = 9 mice for hM4Di^+^. Bar graphs show mean ± SEM. ns = not significant, ** p < 0.01, **** p < 0.0001. Statistical details are shown for significant changes only; all details are provided in statistical details file.

**Fig. S7.**
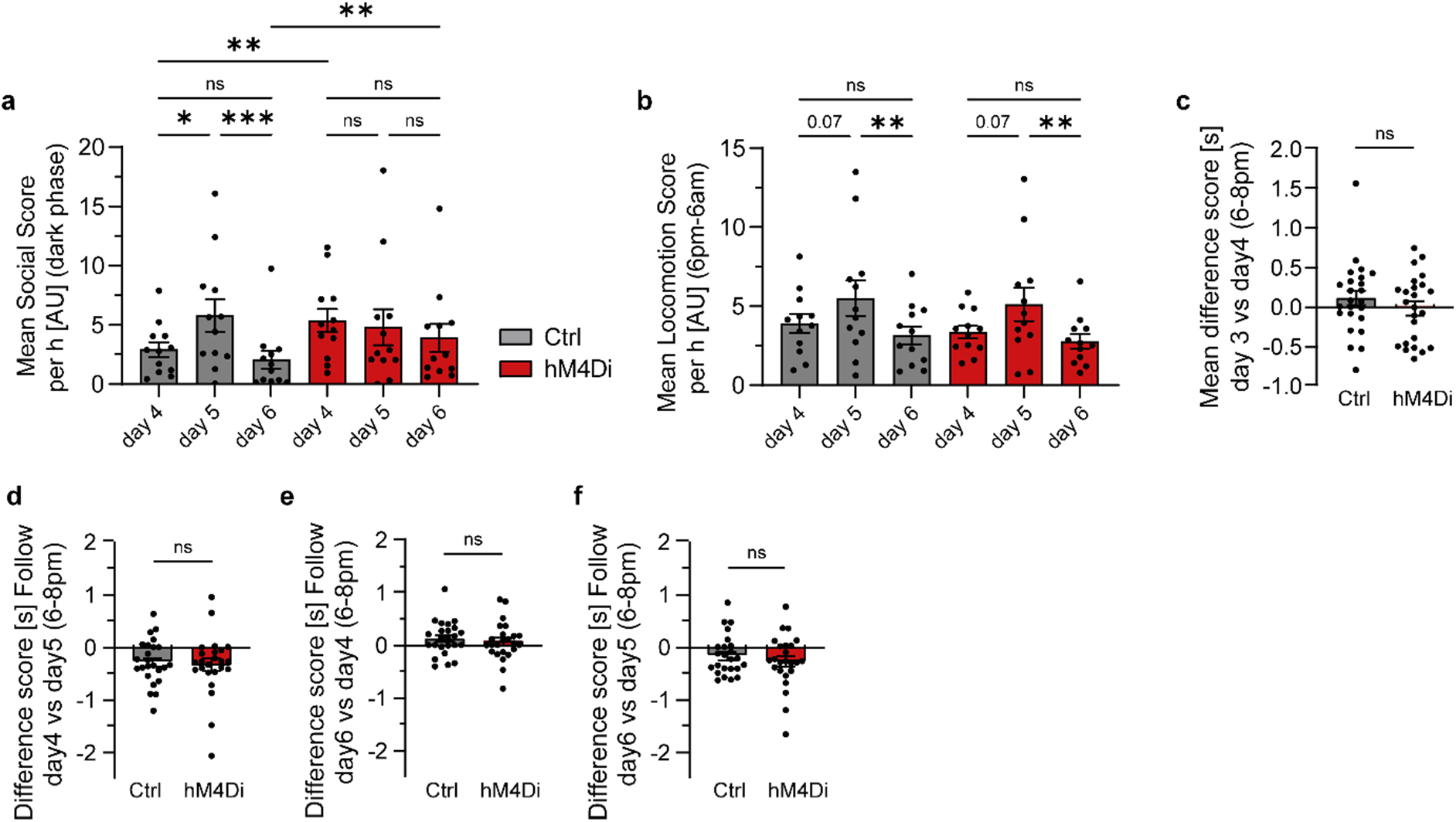
Chronic CA2 inhibition alters social sniffing but not locomotion or social follow behavior of mice. **a** Mean Social Score of groups per hour during dark phase (6pm-6am) on days 4 to 6 of social arena experiments (n = 12h, 2-tailed mixed-effects one-way ANOVA with Geisser-Greenhouse correction and Fisher’s LSD posthoc test). **b** Mean Locomotion Score of groups per hour during dark phase (6pm-6am) on days 4 to 6 of social arena experiments (n = 12h, 2-tailed mixed-effects one-way ANOVA with Geisser-Greenhouse correction and Fisher’s LSD posthoc test). **c** Difference Scores (DS) of social sniffing with familiar mice during the initial 2h of day3 vs day4, calculated in 5min bins. **d-f** Difference Scores (DS) of social follow during the initial 2h during the dark phase, calculated in 5min bins. **d**) DS for interaction with mixed (day 6) vs familiar (day 5) cohort. **e**) DS for interaction with familiar (day4) vs mixed cohort (day5). **f**) DS for interaction with familiar mice before (day4) and after (day6) 24h separation. Statistical details are shown for significant changes only; all details are provided in statistical details file.

